# A Bayesian approach to include Indigenous Knowledge in habitat selection functions

**DOI:** 10.1101/2025.03.26.645387

**Authors:** R Gryba, AL Von Duyke, HP Huntington, B Adams, B Frantz, J Gatten, Q Harcharek, R Sarren, GHR Henry, M Auger-Méthé

## Abstract

Indigenous Knowledge (IK) of species-habitat relationships provides in-depth understanding of species presence. While IK and ‘Western’ scientific data are rarely combined in the same model, Bayesian statistics provides a method to include IK as prior knowledge (priors) and combine it with Western data. In partnership with Iñupiat hunters, we developed a method to include IK in Bayesian habitat selection functions as priors and show how habitat variables mapped by IK holders can be included as covariates. We explain how to include IK and show the effects of IK in a case study for ringed seals (*natchiq* in Iñupiaq; *Pusa hispida*). We show that influence of IK can vary depending on the covariate, and how IK provides information at scales not available in Western data. Our work points to the importance of both Western science and IK data sources in models that may be used for conservation and management decisions.

## I Positionality

This study was initiated after discussions with the North Slope Borough Department of Wildlife Management (DWM), an agency within the regional municipal government, that is dedicated to sustainably managing wildlife populations (including ice associated seals) to ensure that they remain healthy and abundant for subsistence by Iñupiat communities. This study was also reviewed and approved by the Ice Seal Committee (ISC), an Alaskan Native organization “established to help preserve and enhance ice seal habitat; protect and enhance Alaska Native culture, traditions-particularly activities associated with the subsistence use of ice seals” (Ice Seal Committee 2022). This study was developed to meet DWM and ISC mandates to manage ice-associated seals considering both IK and Western scientific knowledge (NSB-DWM 2022; Ice Seal Committee 2022). Co-authors B. Adams, B. Frantz, J. Gatten, Q. Harcharek, and R. Sarren are Iñupiat hunters who reviewed all modelling results to ensure that they align with the Indigenous Knowledge (IK) that they have shared. The other authors are non-Indigenous: R. Gryba was a PhD candidate at the University of British Columbia (UBC), A. Von Duyke is a DWM researcher, M. Auger-Méthé and G. Henry are UBC professors, and H. Huntington is a social scientist. An additional Iñupiaq hunter participated in sharing their IK and providing review but chose to remain anonymous for this publication. Funding for the project was jointly pursued with the DWM, UBC researchers, and with the support of the Native Village of Barrow Iñupiat Traditional Government and ISC. This project was conducted under the UBC Behavioural Research Ethics Board certificate H18-02585.

## 2 Introduction

‘Best available scientific data’ or ‘best available information’ are phrases used to describe data applied to wildlife conservation and management for species and ecosystem protection (e.g., critical habitat designations) (ECCC 2016; NOAA 2023). Under certain jurisdictions, this data or information clearly includes Indigenous Knowledge (IK) (Government of Canada 2002), however, until recently this was not the case (Alexander et al. 2019; Hill et al. 2019; Stern and Humphries 2022). IK can be defined as an “interconnected and systematic knowledge about abiotic and biotic systems and the relationships of those systems with cultural and spiritual aspects of life” (Daniel 2019, pg. 2). Justifications for the omission of IK vary but include a lack of documented IK that could be included within ‘Western’ scientific frameworks, differences in scale between IK and Western science, and/or methodological difficulties with including and combining IK and Western science (e.g., Nadasdy 2003). Two-eyed seeing (Bartlett et al. 2012; Reid et al. 2020), an Indigenous framework that acknowledges the value in different knowledge sources, provides guidance and an opportunity for how to recognize both IK and Western science and address the omission of IK. Although there is a desire to apply ‘two-eyed seeing’ or ‘braiding’ approaches in species management (Stirling et al. 2023), species management remains mostly within colonial frameworks with limited capacity to include different data sources, such as IK, in management decisions (Lamb et al. 2023). As such, there remains a need for methods that can include both IK and Western scientific data into current species management decision-making processes to ensure all of the ‘best available information’ is included.

Efforts to develop methods to include IK as a sole data source in models used to understand species-habitat relationships are ongoing. A common approach has been to use mapped IK as ‘presence-only’ data in species-habitat models. For example, Olsen et al. (2015) used mapped IK of important hunting areas and areas of concentration for bearded seals (*Erignathus barbatus*) as presence data to identify suitable habitat types. Similarly, Evangelista et al. (2018) used IK on species occurrence at specific locations to identify used habitat covariates needed to model and predict the distributions of 38 terrestrial species in Somaliland. Mapped IK of greater bilby (*Macrotis lagotis*) activity was used to identify habitat covariates in Australia to predict greater bilby distribution (Skroblin et al. 2021). Fewer examples exist that use IK of species-habitat relationships directly in modelling. For example, Dubos et al. (2023) used fuzzy logic models to apply IK of Arctic char habitat use (*Salvelinus alpinus*) to model spawning habitat in Nunavut. Gryba et al. (in press) statistically characterized IK of species-habitat relationships to predicted ringed seal (*Pusa hispida*) relative probability of use in Alaskan waters. Polfus et al. (2014) created IK habitat suitability index models for woodland caribou (*Rangifer tarandus caribou*) based on IK species-habitat relationships. While methods that rely on IK as a sole data source to understand species-habitat relationships are valuable in a variety of contexts (e.g., Berkes et al. 2007; Buschman 2022; Stern and Humphries 2022), there is still a need for methods that can include both IK and Western scientific data in the same model. This is particularly important because many current management frameworks still heavily rely on Western scientific data alone and incomplete species life history and habitat use information, which is a hindrance to implementing critical habitat designations (Bird and Hodges 2017).

Bayesian statistics offers a unique framework to model IK and Western scientific data, such as animal movement data, simultaneously. Such modeling frameworks lead to more fully informed models and a better overall understanding of species-habitat relationships. This is possible through the use of ‘informed priors’ that are typically based on previous information such as data collected in previous studies or expert knowledge (Low-Choy et al. 2009). Informed priors describe our current or prior understanding of a system, specifically our existing knowledge of model parameter values (e.g., habitat associations). In most studies, researchers assume that we have no prior information, thus they use priors that are designed to have negligible influence on the final model predictions (often referred to as ‘vague’ priors, Gelman et al. 2014). While Bayesian approaches are designed to be able to use informed priors (Robert 2007), published examples of the use of IK as informed priors are limited, and to our knowledge, none directly include IK of species-habitat relationships in habitat modelling efforts.

There are a few examples of Bayesian analysis with IK as informed priors in related fields, and similarly of the use of non-IK expert knowledge as informed priors in species-habitat models. In a Bayesian time series analysis of sea turtle migratory waves in French Guiana, Girondot and Rizzo (2015) included IK on migration timing as informed priors in the models. In this case, the IK priors allowed for parameter estimation when Western data was limited. IK on polar bears has been collected with the intention of integrating it into Bayesian population models for Alaska polar bears in the Chukchi Sea subpopulation (Braund et al. 2018; Regehr et al. 2018). The few examples where non-IK expert knowledge has been used as informed priors in Bayesian habitat modeling suggest that this approach is promising for the inclusion of IK, a specific type of expert knowledge. For example, Martin et al. (2005) used expert knowledge as priors in models of persistence of multiple bird species in different grazed landscapes. Murray et al. (2009) used informed priors based on the probability of association between rock-wallaby (*Petrogale penicillata*) and habitat covariates based on expert knowledge. Although IK as priors in similar ecological models has been discussed (e.g., Braund et al. 2018), to our knowledge, IK specifically as priors in habitat modeling has not been published.

In this study, we demonstrate the process and value of including IK in Bayesian habitat selection functions (HSFs). We show how to include IK as informed priors and how habitat variables that are mapped by IK holders can be included as a covariate in Bayesian HSFs with satellite telemetry data. To do this we first describe how IK documented in different ways can be statistically characterized to provide point estimates, and in some cases the standard deviation, of the relationships between IK probability of species presence and environmental covariates (based on Gryba et al. in press). We then show how to use the point estimates as IK priors in the HSF and include mapped IK of specific habitat variables in the HSF as a covariate. We demonstrate our approach through a case study of ringed seals in the waters near Utqiaġvik, Alaska in summer; showing how the HSF models that include the IK informed priors and spatial IK covariates differ from those that use vague priors, and result in predicted spatial variation of ringed seal relative probability of use.

## 3 Methods

### 3.1 Habitat Selection Functions

Habitat selection functions (HSFs), sometimes referred to as resource selection functions, are commonly used to quantify habitat selection from an animal’s used locations (e.g., satellite telemetry data; Figure 1.1, 1.2)(e.g., Lele and Keim 2006; Northrup et al. 2022). The goals of HSFs include estimating the covariates’ selection coefficients associated with the selection function. Estimating these coefficients is important because they quantify the strength with which available habitat is selected by the animal (Northrup et al. 2022; Florko et al. in press) and can be used along with environmental covariates to map where the animal is more likely to be.

In practice, HSFs are often applied using a logistic regression, where the used and available data *Y*_*j*_ for *j* = 1, …, *J* locations is assumed to be sampled from a Bernoulli distribution with the probability that the location is used is denoted as *θ*_*j*_. The used locations of *Y*_*j*_ are equal to 1, while the available locations are equal to 0. We use a logit link function to relate the relative probability of the presence of an individual at location *j* to each of the *c* covariates values *x*_*j*1_, …, *x*_*j,c*_ at that location (Fieberg et al. 2021).

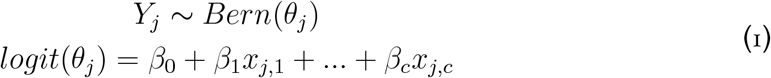

We are primarily interested in estimating *β*_*c*_ for each covariate to understand the relationships between the species and each environmental covariate, and to be able to predict relative probability of use in an area.

HSFs can be fit using a frequentist or Bayesian approach. In a Bayesian framework, typically a vague Gaussian prior would be placed on *β*_*c*_ to allow for the species-habitat relationship to be determined by the data alone,

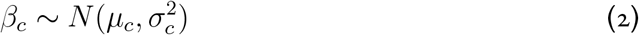

with *µ*_*c*_ = 0 and 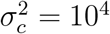.

A Gaussian prior is typically placed on the coefficients because we assume that the coefficient can be any real number and take positive or negative values. One of the benefits of using a Bayesian approach is that informative priors can also be used, adjusting the values of *µ*_*c*_ and 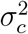 to reflect prior knowledge. The informative prior would then influence the estimated *β*_*c*_ for that coefficient (Robert 2007).

### 3.2 IK as Informed Priors

Indigenous Knowledge specific to species-habitat relationships can be interpreted as a probability of species presence (Gryba et al. in press). Specifically, the proportion of hunters that observed habitat use or habitat relationships by a species can be converted to probability of species presence. For example, if one hunter share that a seal is associated with a specific habitat type, the probability of a seal being associated with that habitat type is set to 1*/n*, where *n* is the number of hunters interviewed. This interpretation allows for the characterization or estimation of the relationships between IK based probability of species presence and environmental covariates. Indigenous Knowledge of species-habitat relationships can be documented in several ways, including quantitative, qualitative, and spatial documentation, each of which can provide information that can be used to inform priors applied to the *β*_*c*_ in HSFs (Figure 1A, B).

Quantitative documentation of IK can be defined as describing IK of a species association with specific, or a range, of habitat values (Gryba et al. in press). For quantitative documentation of IK, a beta regression can be used to estimate the mean response, *µ*_*i*_, of the probability of species presence *Y*_*i*_, here based on the proportion of hunters who shared that information associated with covariate value *i* (Gryba et al. in press):

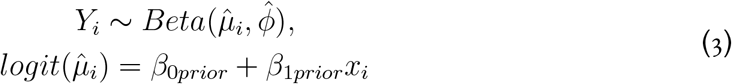

where the *β*_*prior*_ coefficients define the relationship between the IK probability of species presence *Y*_*i*_ and the environmental covariate value *i*. Once this model is fitted to the quantitative IK, the point estimates 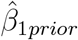 can then be used for the mean of the Gaussian prior distribution *µ*_*c*_ on the regression coefficient *β*_*c*_ in the HSF (Equation 2; James et al. 2010; Kynn 2005). When Bayesian methods are used for the beta regression, the posterior distribution of 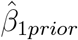 is estimated, and as such the standard deviation, *σ*, of the distribution can also estimated, and applied to the prior distribution.

For qualitative documentation of IK, where the IK holders verbally describe a relationship between a species and habitat characteristics, one must characterize the relationship based on the information provided (Gryba et al. in press). For example, if IK holders describe an increase in species presence with an environmental covariate value, we can use a linear relationship where the slope of the line is calculated based on the IK probability of species presence estimated from the proportion of hunters that shared that IK at the minimum and maximum values of that covariate. The slope can then be used as mean of the Gaussian prior distribution *µ*_*c*_ on the regression coefficient *β*_*c*_ in HSFs (Figure 1E). The 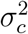 of the IK informed prior is not estimated, and so the researcher, in collaboration with the IK holders, must decide the value to be used based on how much the IK should inform the analysis.

Spatial documentation of IK is shared through maps, where the IK holders identified specific regions with either relatively low or relatively high presence of the animal, or a spatial representation of a habitat variable that may have relatively high or low presence (Gryba et al. in press). The regions can be included in HSFs by creating a raster of the region and assigning a categorical value to each pixel representing whether that pixel is inside or outside of, and sometimes nearby, the region identified by the IK holders. The mean *µ*_*c*_ of the IK informed prior for each category is the IK probability of species presence value, again based on the proportion of hunters that shared that IK. Similar to quantitative documentation of IK, the 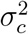 of the IK informed prior is not estimated but can be defined to allow for a highly informative prior or a less informative prior. The spatial documentation IK of habitat variables also provides a good way to include new IK covariates into the HSFs (Figure 1E).

The interpretation and use of IK should only be conducted in partnership with IK holders. This approach only applies a limited portion of what IK is and contains, focused on ecological knowledge within. As such, it is important to recognize the potentially extractive nature of the approach, as these methods and interpretation do not reflect the extent of what IK embodies (Raymond-Yakoubian and Daniel 2018). To ensure continuous involvement and review by IK holders within this method, the statistically characterized IK is mapped and reviewed by the IK holders (Figure 1C, D). If there are any errors or corrections, the process returns to the statistical characterization step (Figure 1B).

**Figure 1.**
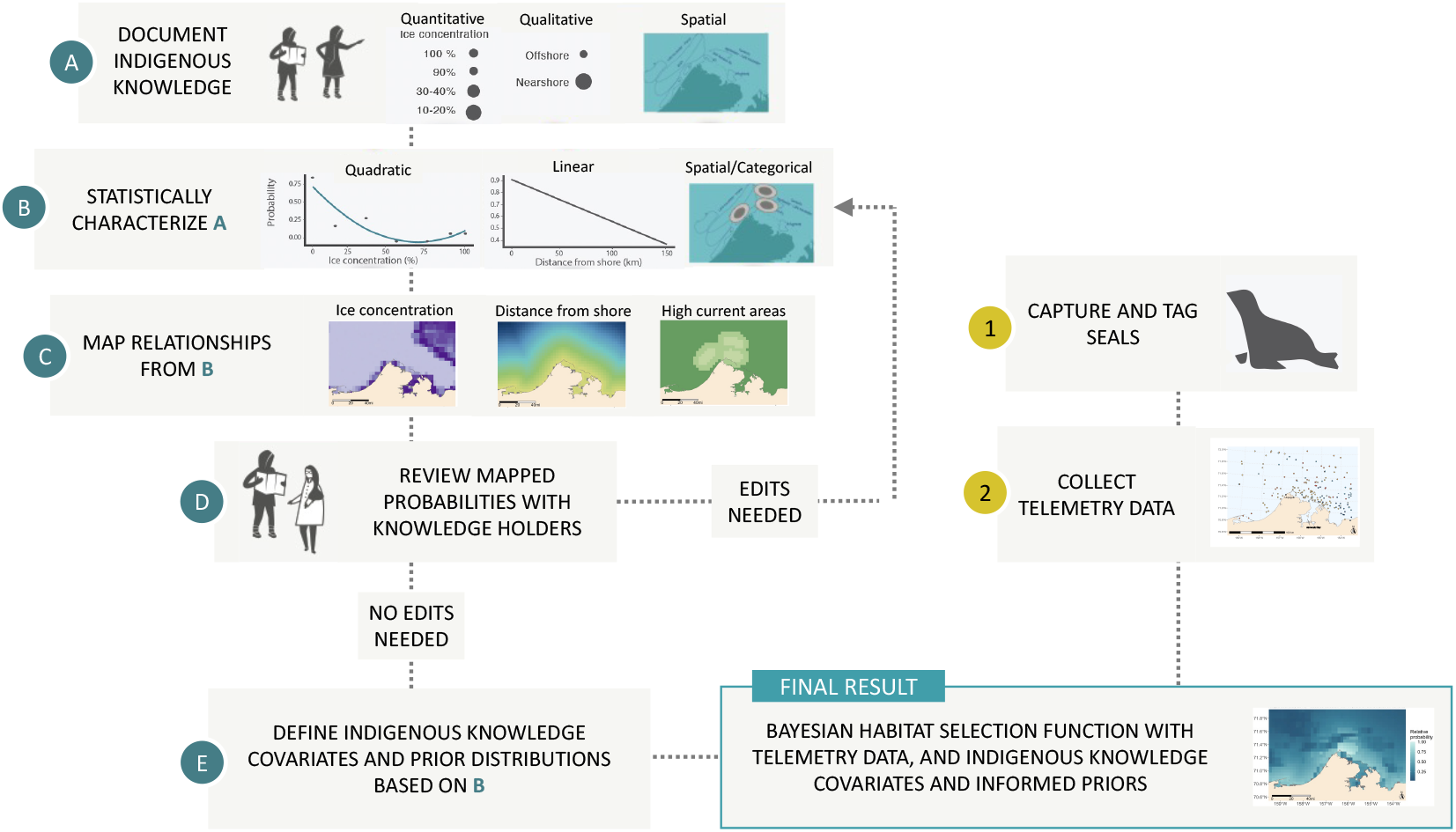
Methods process map to (A) document IK considering three different approaches, (B) statistically characterize the relationships between seals and specific habitat types based on the IK, (C) map the statistically characterized IK of the probability of observing a seal with each habitat type from (B), (D) review the mapped probabilities with the IK holders, retuning to (B) if corrections are needed. Once the statistical characterization is confirmed by the holders, (E) they can be used as covariates and informed priors in HSFs, alongside information from (1) captured and tagged seals and (2) the resulting telemetry data.

### 3.3 Case Study: Ringed Seal IK and Telemetry Data

To illustrate the process of including IK within a statistical framework, we built Bayesian HSF models for ringed seals that included IK as informed priors and as covariates in the model along with satellite telemetry data of ringed seal movement. The IK was documented as part of a collaborative research project with Iñupiat hunters conducted in Utqiaġvik, Alaska, documenting IK of habitat use and behaviour of ice-associated seals, including ringed seals (Gryba et al. 2021). The satellite telemetry data is from ringed seals captured and tagged during the open-water period near Utqiaġvik, Alaska (Figure 2, Table S1) from 2013-2018 (n=13) (VonDuyke et al. 2020).

The locations of the seals included in the analyses were limited to match the spatial extent of the IK shared by hunters from Utqiaġvik, Alaska (Harcharek 2015; Gryba et al. 2021). The location data was temporally limited to July and August - the “summer months” - as defined by IK holders (Gryba et al. 2021).

Seal movements were documented using satellite transmitters (hereafter ‘tags’). Most seals were instrumented with a primary and secondary tag (Wildlife Computers SPLASH and SPOT respectively), the data from which were merged into a single movement timeseries for each seal. In addition to ARGOS locations, the tags recorded dive information (i.e., depth and duration) and behavioural data regarding their haul-out status. The primary tags, which collected most of the data, were attached with glue to the seals’ fur, and were therefore shed during the annual molt the following spring. As such, for each seal, the satellite tag data tended to cover the late summer through early spring of the following year (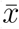 = 229.2 days, range = 39-409 days). The entire data set analyzed ranged from August 10, 2013 to August 31, 2018. ARGOS location data are associated with large measurement errors that are defined by either categorical quality classes or by error ellipses measured along two axes at the location (CLS 2016). Location estimates, considering the ARGOS measurement error, were made using the R-package foieGras (Jonsen and Patterson 2019). Prior to making the location estimates, the first 7 days of locations were removed to account for potential effects from capture and tagging. To meet HSF model assumptions that the data are independent the location data was thinned to one location per day per individual to account for autocorrelation (Figure 2).

**Figure 2.**
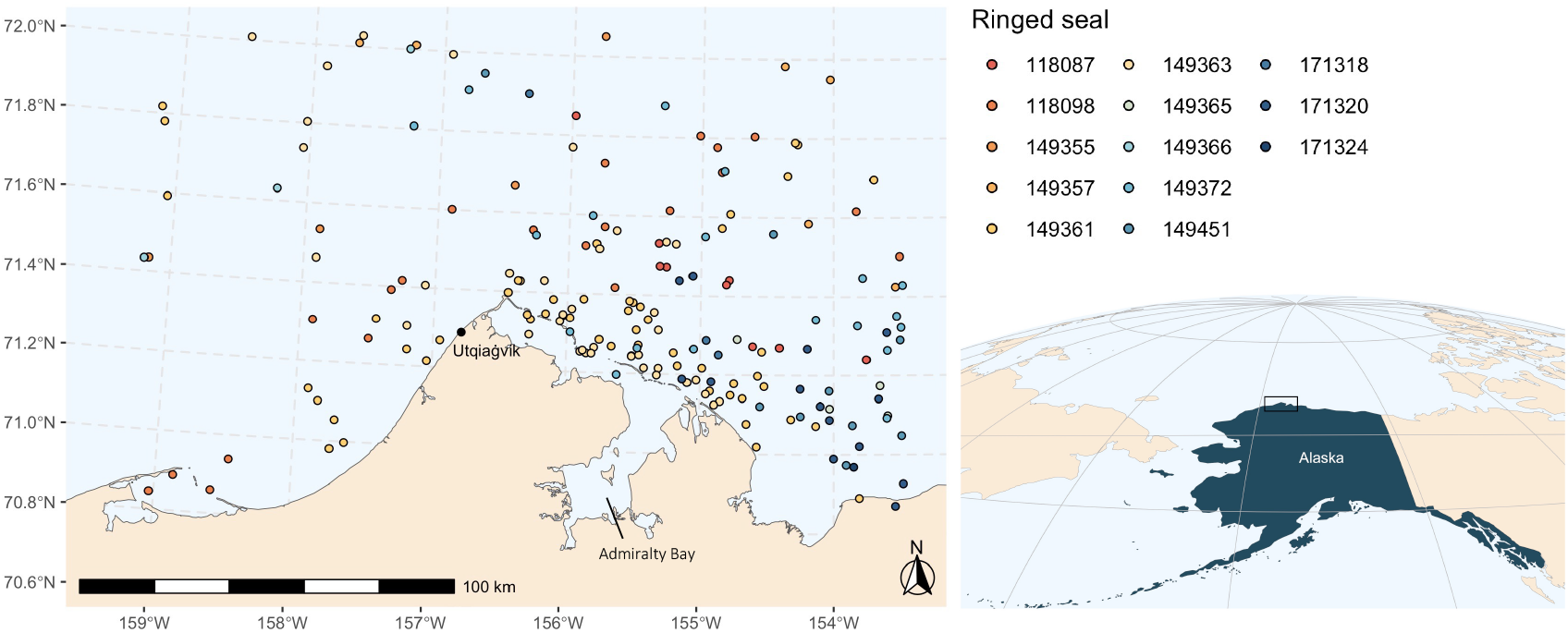
Estimated locations of ringed seals in waters near Utqiaġvik, Alaska July and August, 2013-2018. Colour represents each individual seal.

Habitat suitability functions (equation 1) were applied to the estimated locations. Each HSF included up to four environmental covariates including ice concentration (% ice), distance from shore, currents, and region. These variables were identified by IK holders as important variables that will influence ringed seal habitat selection in summer (Gryba et al. in press). To understand the relative influence of IK within HSF models, we evaluated three HSF model variants: ‘IK-only’, ‘movement-only’, and ‘IK-informed-movement’. Each model variant is detailed below.

#### 3.3.1 IK-only HSF

An ‘IK-only’ HSF was created, as in Gryba et al. (in press), based on the IK of ringed seal habitat use, which was documented as part of an earlier collaborative research project with Iñupiat seal hunters in Utqiaġvik, Alaska (Gryba et al. 2021). Briefly, a beta regression was applied to the quantitative covariate ice concentration (% ice), to estimate *β*_*prior*_ and *σ* associated with the probability of ringed seal presence as related to this covariate (equation 3; Table 1). The beta regression included a quadratic term to reflect a higher probability of ringed seal presence at low and high ice concentrations. A simple decreasing linear relationship was defined for the qualitative IK distance from shore (Table 1). Two spatial covariates were also characterized. Currents that are used by ringed seals in summer were identified by IK holders and then categorized as the main current areas (0-10 km from the currents) and the near current areas (10-20 km from the currents). Each current area was given a probability of species presence based on the proportion of hunters that shared that information (Table 1). Region was dealt with similarly to current areas, with the region outside of Admiralty Bay (Figure 2) given a higher probability of species presence, compared to the area inside Admiralty Bay, again based on the proportion of hunters that shared that information (Table 1). The estimated coefficients 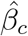 for all covariates were combined in a final equation to calculate the IK relative probability of ringed seal presence. See Gryba et al. (in press) for additional details.

#### 3.3.2 Movement-only HSF

A ‘movement-only’ HSF using the ringed seal satellite telemetry data was run, including distance from shore and ice concentration covariates with vague priors (equation 2). Ice concentration was included as a quadratic in the HSFs to be consistent with the ‘IK-only’ HSF and with IK shared (Gryba et al. in press). The two spatial covariates were not included as they were based on IK and would not typically be included in these models.

#### 3.3.3 IK-informed-movement HSF

An ‘IK-informed-movement’ HSF with the ringed seal satellite telemetry data and IK priors and spatial covariates was run. The IK priors were based on the results from the beta regression and characterization of the quantitative, qualitative and spatial IK used in the ‘IK-only’ HSF (Table 1; Gryba et al. in press). The mean, *µ*_*ice*_, and standard deviation, *σ*_*ice*_, of the IK-prior for ice concentration were the estimated *β*_*prior*_ and *σ* from the beta regression (equation 3; Table 1). The mean, *µ*_*distance*_, of the IK-prior for distance to shore was the slope of the linear relationship (Table 1). Because the IK of distance to shore was qualitative, we had no estimates for *σ*_*distance*_. We therefore chose *σ*_*distance*_ = 1, which increases the importance of the prior in the model, as compared to a vague prior *σ*. To include the currents identified by IK holders as important for ringed seals, a categorical variable was created with three levels: main current areas (0-10 km away), near current areas (10-20 km from the currents), and outside (>20 km). The outside region was the reference level 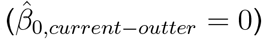.The means, *µ*_*current*−*main*_ and *µ*_*current*−*near*_, for the IK-priors for the other levels were defined using the proportion of hunters that identified these currents as important (Table 1). Similarly, the IK region was also included in the model as a categorical variable, with the area inside Admiralty Bay set as the reference category. Again the IK probability of ringed seal presence associated with the region outside of Admiralty Bay was used for *µ*_*region*_ (Table 1). Since the IK for current areas and the region was spatial, no *σ*_*c*_ was estimated. We selected a value of 1 for all of them (Table 1).

To demonstrate how IK priors change the HSF results, each IK prior was included in the model in turn with a vague prior remaining on the other covariate, and the posteriors were re-estimated. Then each spatial IK covariate was added to the model separately, and again the posteriors were re-estimated, with vague priors on distance from shore and ice concentration. Finally, a model with IK priors used for all the covariates was run.

To estimate the parameters of the HSF, we used the methods outlined in Muff et al. (2020) using R-INLA (Rue et al. 2009; Martins et al. 2013), including weighting of the used and available locations, without random effects. The environmental variables were projected to a 5 km x 5 km grid, Alaska Albers projection, with daily ice concentration from Spreen et al. (2008). Continuous variables were scaled prior to modelling. Available habitat was randomly sampled from the grid with ten times as many samples as those at the used locations, for each date of the used locations. To sample available habitat for ice concentration the date matching the used locations was applied, and the ice concentration in that grid cell for that date was included in the analysis. The same available locations were sampled for the static covariates, ignoring the date.

The resulting HSFs were used to predict ringed seal relative probability of use, with the results scaled by the maximum value. Median predictions were also estimated for 2013-2018 by taking the median value of daily predicted intensity based on the daily ice concentrations, bounded by the satellite telemetry data dates, August 10, 2013-August 31, 2018. The resulting maps were reviewed by the Iñupiat co-authors and an additional hunter who chose to remain anonymous.

**Table 1:** Indigenous Knowledge based *µ*_*c*_ and *σ*_*c*_ for each IK prior. The *µ*_*c*_ values are based on estimates of 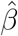 from Gryba et al. (in press) along with the estimate of *σ* for ice concentration. The *σ* values applied for the other covariates were selected to best reflect the IK shared.

| IK type | Covariate | $\mu_c$ | $\sigma_c$ |
| --- | --- | --- | --- |
| Quantitative | Ice concentration | 0.55 | 0.50 |
|  | Ice concentration<sup>2</sup> | 0.87 | 0.77 |
| Qualitative | Distance from shore | -0.16 | 1 |
| Spatial | IK main current area (0-10 km) | 0.11 | 1 |
|  | IK near current area (10-20 km) | 0.08 | 1 |
|  | IK Admiralty Bay region (outside) | 0.89 | 1 |

## 4 Results

The median prediction of relative probability of use for the summer across 2013-2018 showed differences between the ‘IK-only’, ‘IK-informed-movement’, and ‘movement-only’ HSFs, with the combined ‘IK-informed-movement’ model reflecting both data sources (Figure 3A-C). The relative probability of ringed seal presence for the ‘IK-only’ HSF (Figure 3A) showed higher relative probability in the area where currents were identified and in parts of Admiralty Bay, and a decrease in relative probability with distance from shore. The higher association in Admiralty Bay was due to the median values of ice in the area. The median prediction for the ‘movement-only’ HSF with vague priors (Figure 3C), showed a similar decrease in relative probability with distance from shore as the one characterized by the ‘IK-only’ HSF, but did not include nor reflect the relationship ringed seals have with currents. The median prediction for the ‘IK-informed-movement’ HSF with the IK informed priors and covariates (Figure 3B) provided a combined understanding of the relative probability of ringed seal habitat use, reflecting the associations with IK currents, decreased relative probability of use in Admiralty Bay and with distance from shore.

**Figure 3.**
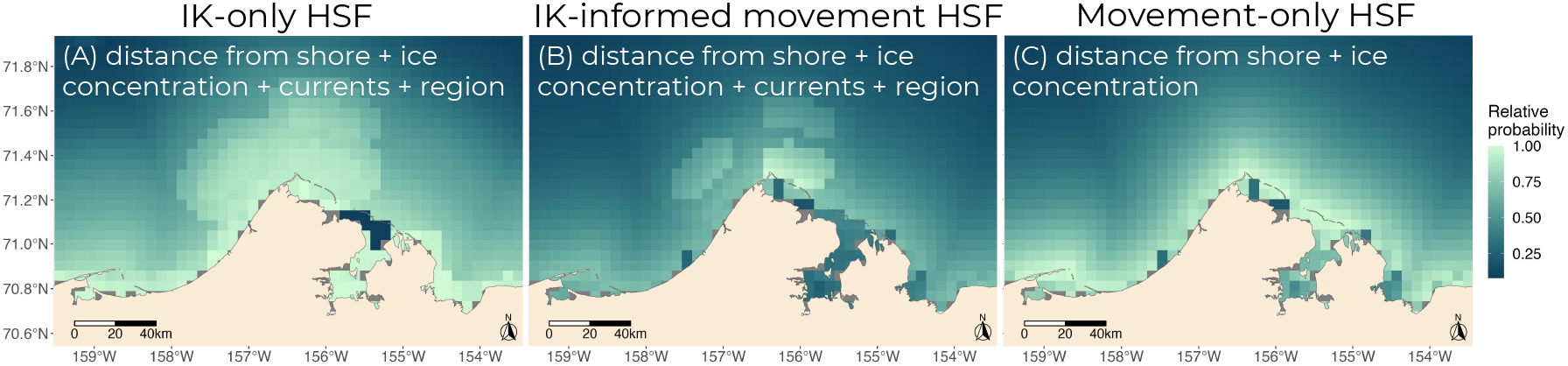
Median predicted summer (2013-2018) relative probability of ringed seal habitat use from (A) the ‘IK-only’ HSF, (B) the ‘IK-informed-movement’ HSF, and (C) the ‘movementonly’ HSF with the variables included in each model.

Adding one IK-informed covariate at a time demonstrated that the inclusion of IK region and current information had the largest impact on the model results (Figure 4). These results are notable as both these covariates are not included in the ‘movement-only’ HSF nor would they have typically been included in the HSF if IK was not considered. For example, the ‘movement-only’ HSF predicted for August 15, 2016 showed a higher relative probability of use closer to shore including in Admiralty Bay (Figure 4A), but adding the IK region to the model as a categorical variable with IK priors (Figure 4B) resulted in a clear decrease in the relative probability of ringed seal presence in Admiralty Bay (Figure 4C). Similarly, when IK currents (Figure 4D) were added as a covariate with an IK prior to the ‘movement-only’ HSF, the relative probability of ringed seal presence increased north of the point (Figure 4E), aligning with the currents identified by the IK holders. In contrast, when the distance from shore IK (Figure 4F) was added as an informed prior to the ‘movement-only’ HSF, there was little change in the prediction (Figure 4G). This lack of change reflected the similarity between the animal movement data and the IK prior, both of which had a negative relationship with distance from shore (Figure S1). This similarity was also apparent in the posteriors for distance from shore between the ‘movement-only’ HSF and the ‘IK-informed-movement’ posterior which overlap (Figure 5). When IK priors for ice concentration (Figure 4H-I) were added to the ‘movement-only’ HSF, there was limited influence of the IK priors, with small difference between the prediction (Figure 4J) and the original ‘movement-only’ HSF (Figure 4A). The effect of the IK priors on the posteriors for ice concentration shifted the linear term from negative to positive, reflecting the higher association with higher ice concentrations noted in the IK, but the IK prior had little influence on the quadratic ice concentration term (Figure 5, Table S2). When there is limited ice present, such as on August 15, 2016 (Figure 4H), the IK prior (Figure 4I) shows higher relative probability of ringed seal presence in the open water areas. This relationship was more complicated when ice was present early in the season in the area.

Relative probability of ringed seal habitat use when ice was present (i.e., July 20, 2016, Figure 6A-C) compared to a day that reflects seasonal norms of open water (i.e., August 15, 2016, Figure 6D-F) showed differences between the ‘IK-only’ HSF, the ‘IK-informed-movement’ HSF, and the ‘movement-only’ HSF results, and the complexity of the relationship with ice concentration. The higher relative probability of ringed seal presence in higher ice concentration areas (Figure 6A, Figure S2) in the ‘IK-only’ HSF reflects the IK that ringed seals will haul out on sheets of ice when present in summer (Gryba et al. 2021; in press). When compared to the predicted relative probability of ringed seal presence from the ‘IK-informed-movement’ HSF, that included all the covariates with IK priors (Figure 6B), and the ‘movement-only’ HSF (Figure 6C), neither reflect the IK association with areas of higher ice concentrations (Figure S2). In contrast, the predictions for an example later in the season, August 20, 2016 (Figure 6D-F), the ‘IK-informed-movement’ HSF showed a combination of the two knowledge sources, with higher relative probability of ringed seals presence in the IK current areas, lower relative probability in Admiralty Bay, and the consistent relationship with distance from shore. Importantly, regardless of ice concentration the ‘IK-only’ and ‘IK-informed-movement’ HSFs both shows higher relative probability in the main current areas and lower relative probability in Admiralty Bay, again reflecting the IK shared (Gryba et al. 2021).

**Figure 4.**
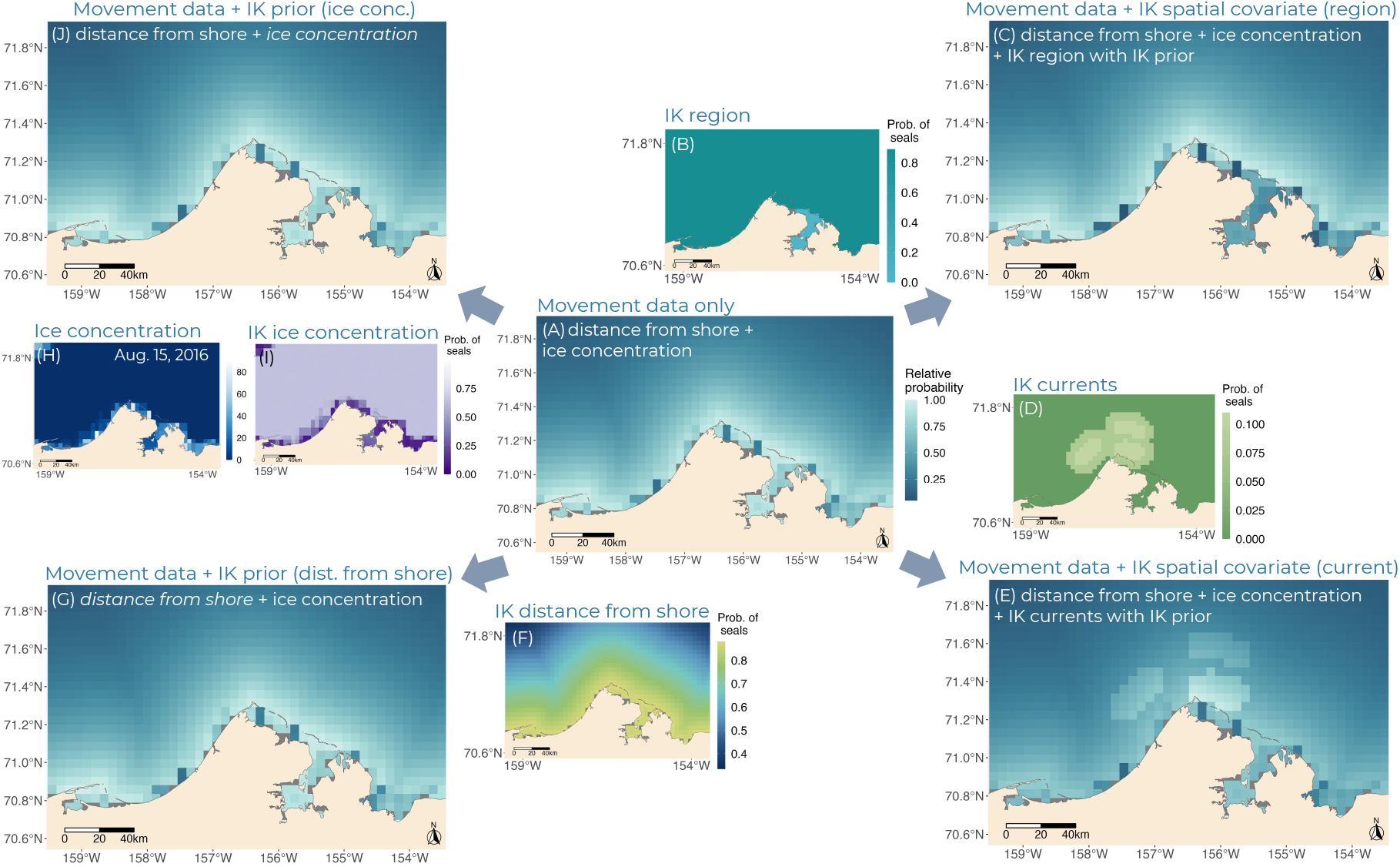
Predicted relative probability of ringed seal presence for August 15, 2016 from the (A) ‘Movement-only’ HSF with distance from shore and ice concentration and vague priors, (B) IK summer probability of ringed seal presence for the region outside of Admiralty Bay, (C) ‘IK-informed-movement’ HSF with distance from shore and ice concentration, and IK region, with IK prior on region, (D) IK summer probability of ringed seal presence for current areas, (E) ‘IK-informed-movement’ HSF with distance from shore and ice concentration, and IK current areas, with an IK prior on currents, (F) IK summer probability of ringed seal presence for distance from shore, (G) ‘IK-informed-movement’ HSF with distance from shore and ice concentration, with an IK prior on distance from shore, (H) Ice concentration (%) on August 15, 2016, (I) IK summer probability of ringed seal presence associated with ice concentration, and (J) ‘IK-informed-movement’ HSF with distance from shore and ice concentration, with IK priors on ice concentration. Predicted relative probability maps - areas of dark blue indicate lower relative probability of ringed seals, light blue indicates areas of higher relative probability. Inset maps showing the IK probability of ringed seal presence are from Gryba et al. (in press).

**Figure 5.**
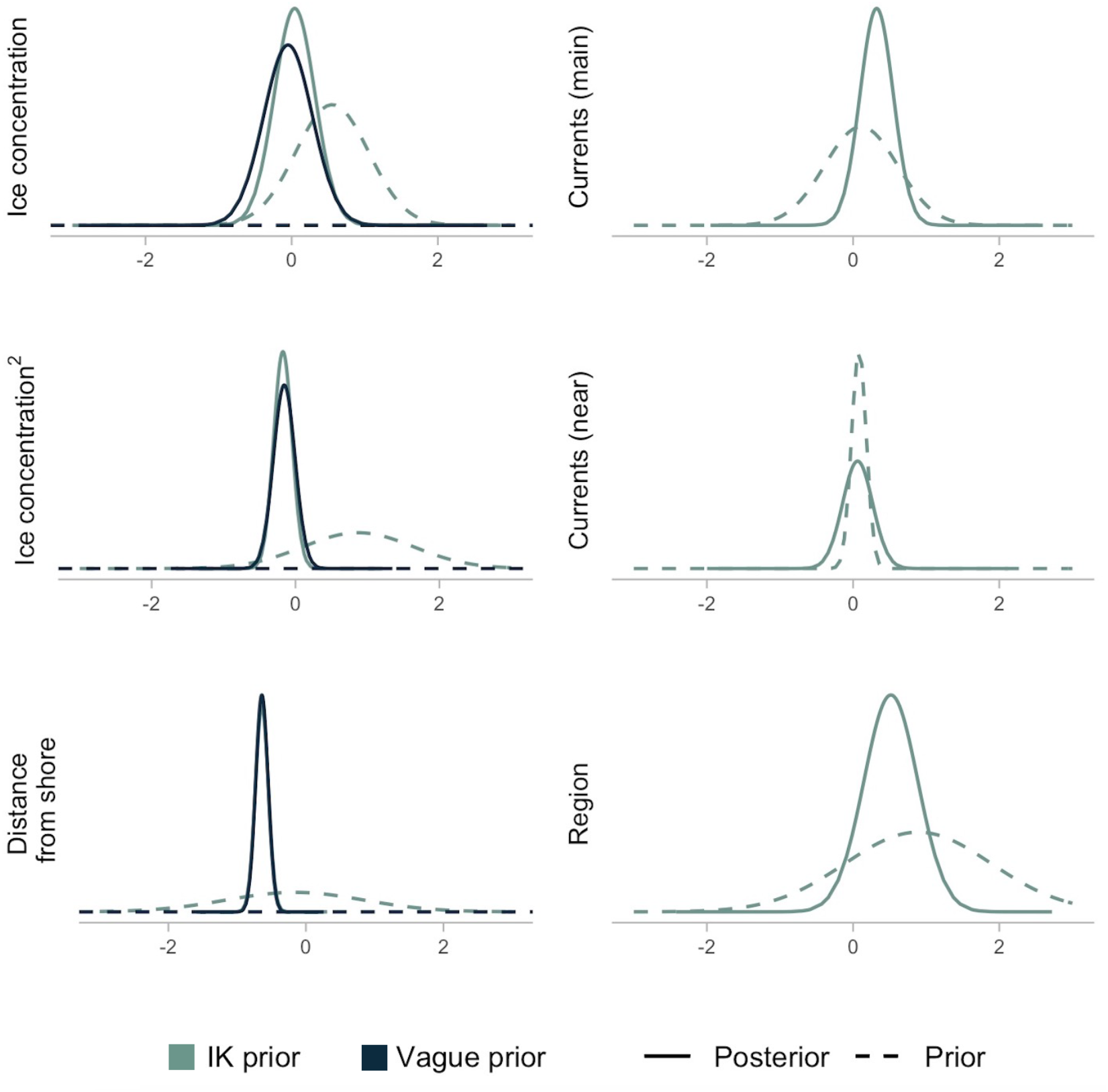
Posteriors and priors for each covariate from the ‘movement-only’ and ‘IK-informed-movement’ HSFs. The results from the ‘movement-only’ HSF with vague priors are shown in dark blue and the ‘IK-informed-movement’ HSF with IK informed priors shown in light blue. The solid lines are the posteriors and the dotted lines are the priors. Region indicates the areas outside of Admiralty Bay, currents (main) is the area at the currents to 10km away, currents (near) is the area from 10-20km away from the currents.

**Figure 6.**
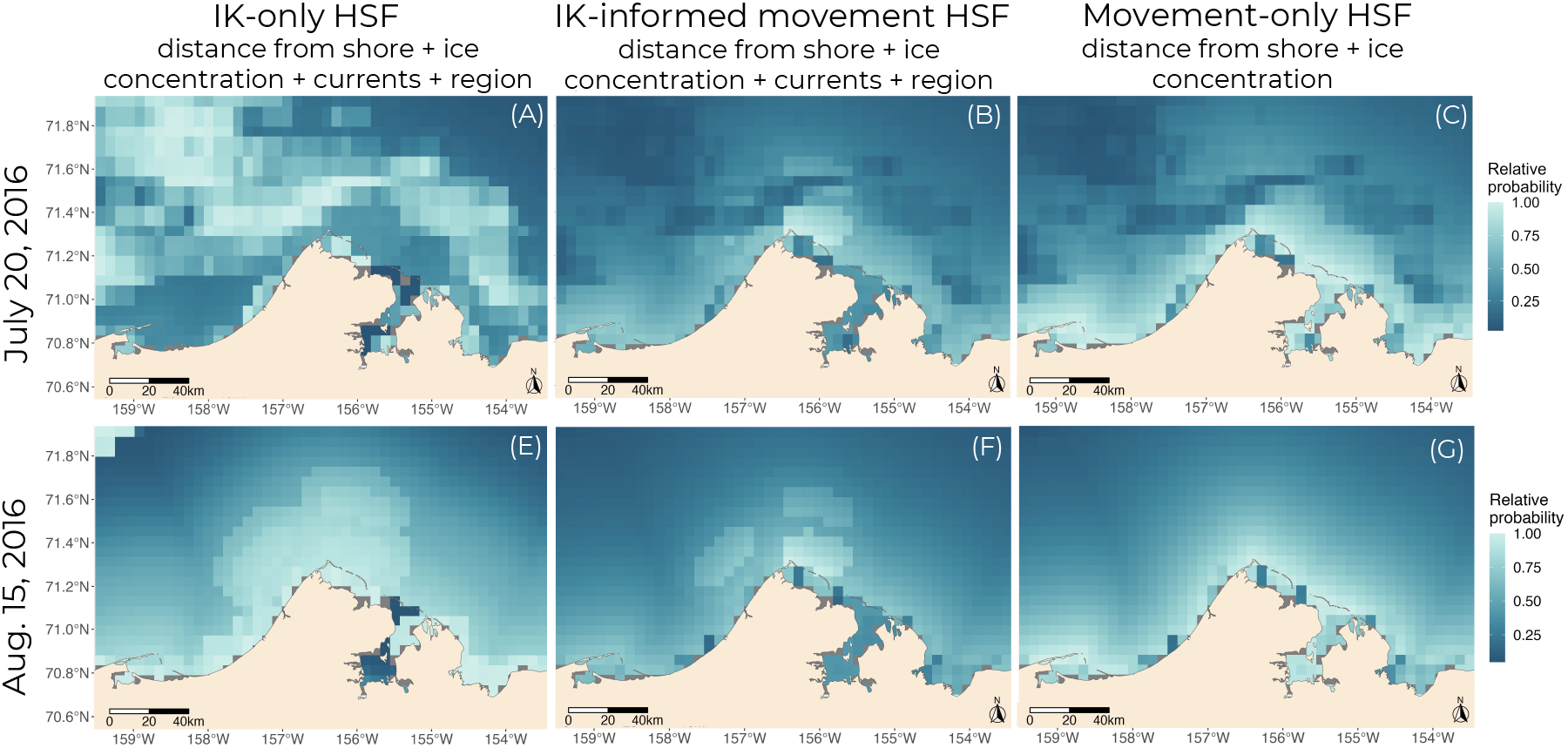
Predicted relative probability of ringed seal use for July 20, 2016 from (A) the ‘IK-only’ HSF including distance from shore, ice concentration, currents, and region, (B) the ‘IK-informed-movement’ HSF including distance from shore, ice concentration, currents, and region, and (C) the ‘movement-only’ HSF with distance from shore and ice concentration with vague priors, and for August 15, 2016 from (D) the ‘IK-only’ HSF including distance from shore, ice concentration, currents, and region, (E) the ‘IK-informed-movement’ HSF including distance from shore, ice concentration, currents, and region, and (F) the ‘movement-only’ HSF with distance from shore and ice concentration with vague priors.

## 5 Discussion

Our study provides a method to include IK as informed priors for habitat relationship coefficients in HSFs. In addition, we show how spatial IK can be included as new categorical variables which allows for the inclusion of environmental covariates that may not be available from other data sources at an appropriate scale. In our case study, the available data on currents (i.e., current direction and speed) in the nearshore waters around Utqiaġvik, Alaska are patchy and coarse in resolution (e.g., 1/12 degree grids; Cassano et al. 2017). Additionally, the resolution of the ARGOS satellite telemetry data available is not fine enough to detect the use of currents by seals (VonDuyke et al. 2020; Gryba et al. 2021). These differences in scale between IK and Western science data have also been noted for other species (e.g., Fraser et al. 2006; Gagnon and Berteaux 2009; Dubos et al. 2023). The inclusion of the in-depth local knowledge of IK holders, related to currents and region, refined the predictions of the relative probability of ringed seal habitat use to reflect information known by IK holders and included environmental covariates that may have otherwise been excluded. In our case study, the IK of distance from shore and the relationships between the satellite telemetry data and distance from shore were very similar. Unsurprisingly, this points to circumstances where these two knowledge sources match. Such similarities will likely be the case for many habitat covariates, as both knowledge sources are reflecting the same ecological processes.

The inclusion of the IK habitat covariates also highlighted important considerations for species conservation and management, and methodological approaches to inform conservation and management decisions. The application of spatial IK can provide additional information on important regions that may be difficult to quantify based on Western or available data sources, providing opportunities to capture complex indices of species presence. Additionally, in our case study we applied a *σ*_*c*_=1, but the practitioner can adjust this value as needed to increase how informative the priors are, with feedback from IK holders to ensure that results reflect IK. The inclusion of spatial IK and the potential to apply IK priors in other models also provides opportunities to include IK when Western data collection may be limited due to considerations of subsistence activities. For example, NOAA aerial surveys for Arctic mammals do not survey within 30 nautical miles of subsistence communities such as Utqiaġvik, Wainwright, Point Lay, Point Hope, and Kivalina, Alaska, to avoid disturbing subsistence species (E. Moreland pers. comm. 2023). The application of spatial IK and IK priors in this context could provide opportunities to supplement information on the relative probability of species habitat use. For other species, and for other regions, IK could similarly identify areas with increased species presence not reflected in Western science data sources, and the inclusion of IK areas or regions as categorical variables provides a simple solution for their inclusion as covariates in HSFs.

The inclusion of IK as informed priors in HSFs had differing influences on the results based on how informative the priors were but also because of how the satellite telemetry data was collected. In our case study, the minimal influence of the IK priors on the estimated 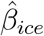 and 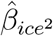 is largely a result of when ringed seals are tagged in this area, the ice conditions required for capture and tagging, and as a result the satellite telemetry data available. Typically, ringed seals are captured and tagged near Utqiaġvik, Alaska when the ice has started to break up and open water areas are present to be able to set nets (VonDuyke et al. 2020). Based on capture timing and the removal of the first seven days of data in an attempt to address potential behavioural response to capture and tagging, the majority of seal locations were associate with ice concentrations well below 50% (Figure S3). Additionally, because of the season, the majority of the area was ice free for the rest of the summer. IK holders indicated that when ice sheets are present, they will see ringed seals hauled out on that ice, but if there is no ice present, then ringed seals will be observed throughout the open water areas but with higher relative probability of presence in near shore areas, with currents, and outside of Admiralty Bay (Gryba et al. 2021). Although the predictions from the ‘IK-informed-movement’ HSF for earlier in the season, when higher ice concentrations are present, do not reflect IK, the predictions during open water, that overlap with the satellite telemetry data temporally, also accurately reflect IK during that time. This is an important consideration when interpreting seasonal HSFs and also when including IK and interpreting what may be initially, and erroneously, considered a mismatch between the two knowledge sources. In our case study, it is possible, and seems likely, that the satellite data is not capturing all aspects of habitat use by ringed seals in this region, where as the IK provides an extended insight into broad habitat use that can occur throughout the summer. Similar to Gagnon and Berteaux (2009), we found when comparing the IK to the available Western science data, the IK provided an expanded view into ringed seal habitat use. This points to the importance of both data sources and consideration of how and when Western science data is collected and the information it is actually providing on species habitat use. It is important to note that including IK as priors does not equally weigh the IK and the movement data, as large amounts of data will dominate the results (Robert 2007). It is possible to increase how informative the IK priors are, as noted, but other methods such as including IK as a joint likelihood in the model could also be explored.

The inclusion of IK into HSFs specific to ringed seals are important in the context of the current ringed seal critical habitat designation in US waters (NOAA 2022). The current designated critical habitat does not include the nearshore areas along the Alaska coast (NOAA 2022). The reason provided is the requirement of water depths *>* 2 m when considering habitat requirements for building subnivean birthing lairs and to avoid areas where bottom-fast ice is likely to occur (NOAA 2022). The importance of nearshore habitat to ringed seals, highlighted here in the satellite telemetry data, the IK, and the resulting HSF, was also reflected in comments made by IK holders during the review of the critical habitat designation who pointed to the importance of nearshore habitat in areas such as Seward Peninsula, Norton Sound, and Kotzebue Sound (NOAA 2022). The inclusion of IK directly in policy decisions increases the reliability of a system when diverse knowledge is included and considered, and resulting decisions are also more reliable when they consider all available information (Alexander et al. 2019). As noted by Taqulik Hepa, Director of the NSB-DWM and an Iñupiaq subsistence hunter, “Satellite data can show, for example, a void spatially, but asking hunters about those areas provides different information. There are similarities between satellite tags and IK but satellite tags can miss things.”

Although the inclusion of IK in species-habitat modelling has increased in recent years, the use of IK as Bayesian informed priors has had limited applications. While other published studies have importantly shown how IK can be used to indirectly provide information on species-habitat relationships based on IK occurrence data (e.g., Skroblin et al. 2021; Evangelista et al. 2018) our statistical characterization of the relationship between IK probability of species presence and environmental covariates allows for the direct use of IK to define informed priors. Girondot and Rizzo (2015) are one of the few examples where IK was included as an informed prior, with values of the prior distribution of marine turtle nesting timing defined by the IK. Similar to our approach, Girondot and Rizzo (2015) used the IK of nesting month to define the centre of a uniform prior distribution or the mean of a Gaussian prior distribution. The application of expert opinion as Bayesian informed priors by Martin et al. (2005) was similar to our approach, as expert knowledge informed the distribution of a Gaussian prior using the weighted mean of expert scored on the effects of grazing on species abundance. Although these two approaches define the prior distribution based on IK or expert knowledge, our approach differs in application (i.e., use in habitat models) and also relies on the statistical characterization of a species-habitat relationship.

Our methods have provided an approach to include IK as informed priors and spatial covariates in HSFs, and also guidance on the general conversion of IK into informed priors that could be applied in other types of models. We provide examples for quantitative, qualitative, and spatial IK and through a case study show the importance of the inclusion of IK to provide fully informed models that include data that may not be otherwise available (e.g., nearshore currents). The influence of IK as informed priors or as spatial covariates can have varying effects depending on the covariate and highlights the importance of considering when and how Western data sources are collected and how IK can provide additional insights into species habitat use. Our case study also points to the importance of including IK in ringed seal HSFs and shows the value of both Western science and IK data sources when predicting important areas for species that may be relied upon for conservation and management decisions.

## 6 Acknowledgments

Many thanks to the additional hunter who requested to remain anonymous for sharing their knowledge and reviewing all results. Many thanks to Taqulik Hepa for her thoughts, input, and continued support of this project. Funding: This work was funded by the North Pacific Research Board (Project 1815), and supported in part by funding from the Social Sciences and Humanities Research Council Doctoral Fellowship, University of British Columbia Four Year Doctoral Fellowship, ACUNS Dr. Jim McDonald Scholarship, Lawrence Edward Hassell Graduate Field Research Award in Fisheries, Northern Scientific Training Program, Stantec Research & Development funds, Canadian Research Chairs program, Natural Sciences and Engineering Research Council of Canada, BC Knowledge Development fund, and Canada Foundation for Innovation’s John R. Evans Leaders Fund. Ringed seal tagging was conducted under NMFS Research Permits #358-1787 and #15324 and approved by Alaska Department of Fish and Game (ADFG) Animal Care and Use Committee Protocols #2010-13R and #2016-23. This project was conducted under the University of British Columbia Animal Care Committee Certificate A19-0163. Funding for the ringed seal SPLASH and SPOT satellite tags was provided by the North Slope Borough (NSB)-Shell Baseline Studies Program.

## References

Alexander, S.M., Provencher, J.F., Henri, D.A., Taylor, J.J., Lloren, J.I., Nanayakkara, L., Johnson, J.T., & Cooke, S.J. (2019). Bridging Indigenous and science-based knowledge in coastal and marine research, monitoring, and management in Canada. Environmental Evidence, 8:36.

Bartlett, C., Marshall, M., & Marshall, A. (2012). Two-eyed seeing and other lessons learned within a co-learning journey of bringing together Indigenous and mainstream knowledges and ways of knowing. Journal of Environmental Studies and Sciences, 2(4):331–340.

Berkes, F., Berkes, M.K., & Fast, H. (2007). Collaborative integrated management in Canada’s North: The role of local and traditional knowledge and community-based monitoring. Coastal Management, 35(1):143–162. doi: 10.1080/08920750600970487.

Bird, S.C. & Hodges, K.E. (2017). Critical habitat designation for Canadian listed species: Slow, biased, and incomplete. Environmental Science & Policy, 71:1–8. doi: 10.1016/j.envsci.2017.01.007.

Braund, S.R., Lawrence, P.B., Sears, E.G., Schraer, R.K., Regehr, E.V., Adams, B., Hepa, R.T., George, J.C., & Duyke, A.L.V., (2018). Polar Bear TEK: A pilot study to inform polar bear management models. Technical report, North Slope Borough Department of Wildlife

Management, North Slope Borough Department of Wildlife Management, Research Report NSB.DWM.RR.2018-01. Utqiag?vik, Alaska USA. Available at: www.north-slope.org/departments/wildlife-management. Issue: NSB.DWM.RR.2018-01.

Buschman, V.Q. (2022). Framing co-productive conservation in partnership with Arctic Indigenous peoples. Conservation Biology, 00, e13972.

Cassano, J.J., DuVivier, A., Roberts, A., Hughes, M., Seefeldt, M., Brunke, M., Craig, A., Fisel, B., Gutowski, W., Hamman, J., Higgins, M., Maslowski, W., Nijssen, B., Osinski, R., & Zeng, X. (2017). Development of the Regional Arctic System Model (RASM): Near-surface atmospheric climate sensitivity. Journal of Climate, 30:5729–5733.

CLS, (2016). Collecte Localisation Satellites - argos user’s manual. Technical report.

Daniel, R. (2019). Understanding Our Environment Requires an Indigenous Worldview. Eos, 100. doi: 10.1029/2019EO137482.

Dubos, V., St-Hilaire, A., & Bergeron, N.E. (2023). Fuzzy logic modelling of anadromous Arctic char spawning habitat from Nunavik Inuit knowledge. Ecological Modelling, 477:110262. doi: 10.1016/j.ecolmodel.2022.110262.

ECCC. (2016). Critical Habitat Identification Toolbox: Species at Risk Act guidance. Available at: https://www.canada.ca/en/environment-climatechange/services/species-risk-public-registry/critical-habitat-descriptions/identification-toolbox-guidance.html.

Evangelista, P.H., Mohamed, A.M., Hussein, I.A., Saied, A.H., Mohammed, A.H., & Young, N.E. (2018). Integrating indigenous local knowledge and species distribution modeling to detect wildlife in Somaliland. Ecosphere, 9(3):e02134. doi: 10.1002/ecs2.2134.

Fieberg, J., Signer, J., Smith, B., & Avgar, T. (2021). A ‘How to’ guide for interpreting parameters in habitat-selection analyses. Journal of Animal Ecology, 90(5):1027–1043. doi: 10.1111/1365-2656.13441.

Florko, K.R.N., Togunov, R.R., Gryba, R., Sidrow, E., Ferguson, S.H., Yurkowski, D.J., & Auger-Méthé, M. (in press). An introduction to statistical models used to characterize species-habitat associations with animal movement data. Movement Ecology.

Fraser, D.J., Coon, T., Prince, M.R., Dion, R., & Bernatchez, L. (2006). Integrating Traditional and Evolutionary Knowledge in Biodiversity Conservation: a Population Level Case Study. Ecology and Society, 11(2). doi: 10.5751/es-01754-110204.

Gagnon, C.A. & Berteaux, D. (2009). Integrating Traditional Ecological Knowledge and ecological science: a question of scale. Ecology and Society, 14(2):19. doi: 10.5751/ES-02923-140219. Publisher: The Resilience Alliance.

Gelman, A., Carlin, J.B., Stern, H.S., & Rubin, D.B., (2014). Bayesian data analysis. Chapman & Hall/CRC Press, Boca Raton, FL, USA.

Girondot, M. & Rizzo, A. (2015). Bayesian Framework to Integrate Traditional Ecological Knowledge into Ecological Modeling: A Case Study. Journal of Ethnobiology, 35(2):337–353. doi: 10.2993/etbi-35-02-337-353.1.

Government of Canada. (2002). Species at risk act (s.c. 2002, c. 29). Available at: https://laws-lois.justice.gc.ca/eng/acts/s-15.3/page-2.html.

Gryba, R., Huntington, H., Von Duyke, A., Adams, B., Frantz, B., Gatten, J., Harcharek, Q., Olemaun, H., Sarren, R., Skin, J., Henry, G., & Auger-Méthé, M. (2021). Indigenous Knowledge of bearded seal (Erignathus barbatus), ringed seal (Pusa hispida), and spotted seal (Phoca largha) behaviour and habitat use near Utqiag?vik, Alaska, USA. Arctic Science, 7(4):832–858. doi: 10.1139/as-2020-0052.

Gryba, R., VonDuyke, A., Huntington, H., Adams, B., Frantz, B., Gatten, J., Harcharek, Q., Sarren, R., Henry, G., & Auger-Méthé, M. (in press). Indigenous Knowledge as a sole data source in habitat selection functionse. PNAS. doi: 10.1101/2023.09.07.556613.

Harcharek, Q., (2015). Spatial analysis of subsistence with GPS. Technical report, North Slope Borough, North Slope Borough. Barrow, AK.

Hill, C.J., Schuster, R., & Bennett, J.R. (2019). Indigenous involvement in the Canadian species at risk recovery process. Environmental Science & Policy, 94:220–226. doi: 10.1016/j.envsci.2019.01.017.

Ice Seal Committee. (2022). Ice Seal Committee: Co-Management of Alaska’s Ice Seals. Available at: https://www.iceseals.org/.

James, A., Choy, S.L., & Mengersen, K. (2010). Elicitator: An expert elicitation tool for regression in ecology. Environmental Modelling and Software, 25(1):129–145. doi: 10.1016/j.envsoft.2009.07.003. Publisher: Elsevier Ltd.

Jonsen, I. & Patterson, T.A. (2019). foieGras: Fit continuous-time state-space and latent variable models for filtering Argos satellite (and other) telemetry data and estimating movement behaviour.

Kynn, M., (2005). Eliciting expert knowledge for Bayesian logistic regression in species habitat modelling. PhD thesis, Queensland University of Technology.

Lamb, C.T., Willson, R., Menzies, A.K., Owens-Beek, N., Price, M., McNay, S., Otto, S.P., Hessami, M., Popp, J.N., Hebblewhite, M., & Ford, A.T. (2023). Braiding Indigenous rights and endangered species law. Science, 380(6646):694–696. doi: 10.1126/science.adg9830.

Lele, S.R. & Keim, J.L. (2006). Weighted distributions and estimation of resource selection probability functions. Ecology, 87(12):3021–3028.

Low-Choy, S., O’Leary, R., & Mengersen, K. (2009). Eliciation by design in ecology: using expert opinion to inform priors for Bayesian statistical models. Ecology, 90(1):265–277.

Martin, T.G., Kuhnert, P.M., Mengessen, K., & Possingham, H.P. (2005). The power of expert opinion in ecological models using Bayesian methods: Impact of grazing on birds. Ecological Applications, 15(1):266–280. doi: 10.1890/03-5400.

Martins, T.G., Simpson, D., Lindgren, F., & Rue, H. (2013). Bayesian computing with INLA: New features. Computational Statistics & Data Analysis, 67:68–83. doi: 10.1016/j.csda.2013.04.014.

Muff, S., Signer, J., & Fieberg, J. (2020). Accounting for individual-specific variation in habitat-selection studies: Efficient estimation of mixed-effects models using Bayesian or frequentist computation. Journal of Animal Ecology, 89(1). doi: 10.1111/1365-2656.13087.

Murray, J.V., Goldizen, A.W., O’Leary, R.A., McAlpine, C.A., Possingham, H.P., & Choy, S.L. (2009). How useful is expert opinion for predicting the distribution of a species within and beyond the region of expertise? A case study using brush-tailed rock-wallabies Petrogale penicillata. Journal of Applied Ecology, 46(4):842–851. doi: 10.1111/j.1365-2664.2009.01671.x.

Nadasdy, P. (2003). Reevaluating the Co-management Success Story. ARCTIC, 56(4):367–380. doi: 10.14430/arctic634.

NOAA. (2022). 87 FR 19232: Endangered and Threatened Species; Designation of Critical Habitat for the Arctic Subspecies of the Ringed Seal.

NOAA. (2023). Endangered Species Conservation: Critical Habitat. Available at: https://www.fisheries.noaa.gov/national/endangered-species-conservation/critical-habitat.

Northrup, J.M., Vander Wal, E., Bonar, M., Fieberg, J., Laforge, M.P., Leclerc, M., Prokopenko, C.M., & Gerber, B.D. (2022). Conceptual and methodological advances in habitat-selection modeling: guidelines for ecology and evolution. Ecological Applications, 32(1). doi: 10.1002/eap.2470.

NSB-DWM. (2022). North Slope Borough Department of Wildlife Management. Available at: https://ns.texrus.com/departments/wildlife-management/.

Olsen, P.M., Kolden, C.A., & Gadamus, L. (2015). Developing theoretical marine habitat suitability models from remotely-sensed data and traditional ecological knowledge. Remote Sensing, 7(9):11863–11886. doi: 10.3390/rs70911863. ISBN: 1186311886.

Polfus, J.L., Heinemeyer, K., & Hebblewhite, M. (2014). Comparing traditional ecological knowledge and western science woodland caribou habitat models. Journal of Wildlife Management, 78(1):112–121. doi: 10.1002/jwmg.643.

Raymond-Yakoubian, J. & Daniel, R. (2018). An Indigenous approach to ocean planning and policy in the Bering Strait region of Alaska. Marine Policy, 97:101–108. doi: 10.1016/j.marpol.2018.08.028.

Regehr, E.V., Hostetter, N.J., Wilson, R.R., Rode, K.D., Martin, M.S., & Converse, S.J. (2018). Integrated Population Modeling Provides the First Empirical Estimates of Vital Rates and Abundance for Polar Bears in the Chukchi Sea. Scientific Reports, 8(1):16780. doi: 10.1038/s41598-018-34824-7.

Reid, A.J., Eckert, L.E., Lane, J.F., Young, N., Hinch, S.G., Darimont, C.T., Cooke, S.J., Ban, N.C., & Marshall, A. (2020). “Two-Eyed Seeing”: An Indigenous framework to transform fisheries research and management. Fish and Fisheries, 22(April 2020):243–261. doi: 10.1111/faf.12516.

Robert, C.P., (2007). The Bayesian Choice: From Decision-Theoretic Foundations to Computational Implementation. Springer, New York, New York.

Rue, H., Martino, S., & Chopin, N. (2009). Approximate Bayesian inference for latent Gaussian models by using integrated nested Laplace approximations. Journal of the Royal Statistical Society: Series B (Statistical Methodology), 71(2):319–392. doi: 10.1111/j.1467-9868.2008.00700.x.

Skroblin, A., Carboon, T., Bidu, G., Chapman, N., Miller, M., Taylor, K., Taylor, W., Game, E.T., & Wintle, B.A. (2021). Including indigenous knowledge in species distribution modeling for increased ecological insights. Conservation Biology, 35(2):587–597. doi: 10.1111/cobi.13373.

Spreen, G., Kaleschke, L., & Heygster, G. (2008). Sea ice remote sensing using AMSR-E 89 GHz channels. Journal of Geophyical Research: Oceans, 113(C2):1–14.

Stern, E.R. & Humphries, M.M. (2022). Interweaving local, expert, and Indigenous knowledge into quantitative wildlife analyses: A systematic review. Biological Conservation, 266:109444. doi: 10.1016/j.biocon.2021.109444.

Stirling, K.M., Almack, K., Boucher, N., Duncan, A., Muir, A.M., Connoy, J.W., Gagnon, V.S., Lauzon, R.J., Mussett, K.J., Nonkes, C., Vojno, N., & Reid, A.J. (2023). Experiences and insights on Bridging Knowledge Systems between Indigenous and non-Indigenous partners: Learnings from the Laurentian Great Lakes. Journal of Great Lakes Research, 49:S58–S71. doi: 10.1016/j.jglr.2023.01.007.

VonDuyke, A.L., Douglas, D.C., Herreman, J., & Crawford, J.A. (2020). Ringed seal (Pusa hispida) spatial use, dives, and haul-out behavior in the Beaufort, Chukchi, and Bering Seas (2011-2017). Ecology and Evolution, 00:1–22. doi: 10.13140/RG.2.2.29160.47368.

