## Supplementary Material for "A Bayesian approach to include Indigenous Knowledge in habitat selection functions"

### 1 Supplementary Material

Table S1: Ringed seals captured and tagged near Utqiaġvik, Alaska, 2013-2018, included in analysis. F = Female, M = Male, Juv = Juvenile. Life stage was determined based on the number of claw bands.

| | ID | Date tagged | Last tag transmission date | Sex | Life stage |
|---|---|---|---|---|---|
| 1 | 118087 | 2013-07-17 | 2013-08-25 | F | Adult |
| 2 | 118098 | 2014-07-22 | 2015-07-12 | M | Juv |
| 3 | 149357 | 2016-07-01 | 2017-02-12 | M | Adult |
| 4 | 149371 | 2016-07-01 | 2017-01-28 | F | Juv |
| 5 | 149451 | 2016-07-01 | 2017-03-24 | M | Adult |
| 6 | 149366 | 2016-07-01 | 2017-01-24 | F | Juv |
| 7 | 149363 | 2016-07-02 | 2017-03-02 | F | Adult |
| 8 | 149355 | 2016-07-02 | 2017-02-11 | F | Juv |
| 9 | 149361 | 2016-07-02 | 2017-02-27 | M | Adult |
| 10 | 149365 | 2016-07-02 | 2017-01-08 | M | Adult |
| 11 | 171324 | 2017-07-03 | 2018-08-16 | M | Adult |
| 12 | 171320 | 2018-07-09 | 2019-02-21 | F | Juv |
| 13 | 171318 | 2018-07-10 | 2018-11-29 | M | Adult |

Table S2: Habitat selection posterior estimates of $\beta_c$ and $\sigma_c$ for the ringed seal 'movement-only' HSF that included vague priors and the 'IK-informed-movement' HSF for ice concentration (ice, ice$^2$), distance from shore (dist), IK current areas ('main' for in the currents identified and up to 10 km away, 'near' for the area 10-20 km from the currents), and IK region (for the area outside of Admiralty Bay). Currents and region were only included in the 'IK-informed' HSF.

| Parameter | Model with vague priors | Model with IK priors |
|---|---|---|
| $\beta_{ice}$ | -0.05 | 0.04 |
| $\sigma_{ice}$ | 0.33 | 0.28 |
| $\beta_{ice^2}$ | -0.16 | -0.18 |
| $\sigma_{ice^2}$ | 0.15 | 0.13 |
| $\beta_{dist}$ | -0.64 | -0.64 |
| $\sigma_{dist}$ | 0.09 | 0.10 |
| $\beta_{currents-main}$ | - | 0.32 |
| $\sigma_{currents-main}$ | - | 0.23 |
| $\beta_{currents-near}$ | - | 0.06 |
| $\sigma_{currents-near}$ | - | 0.21 |
| $\beta_{region}$ | - | 0.52 |
| $\sigma_{region}$ | - | 0.37 |

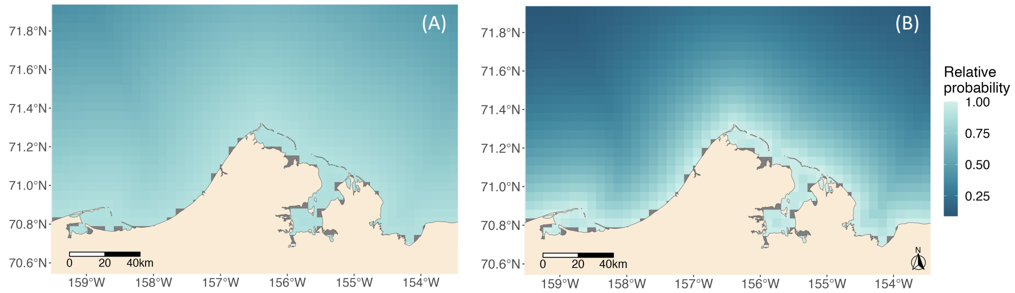


Figure S1: Relative predicted probability of ringed seal presence in the summer from: (A) 'IK-only' HSF model with only distance from shore as a covariate, (B) 'movement-only' HSF with distance from shore as the only covariate with a vague prior.

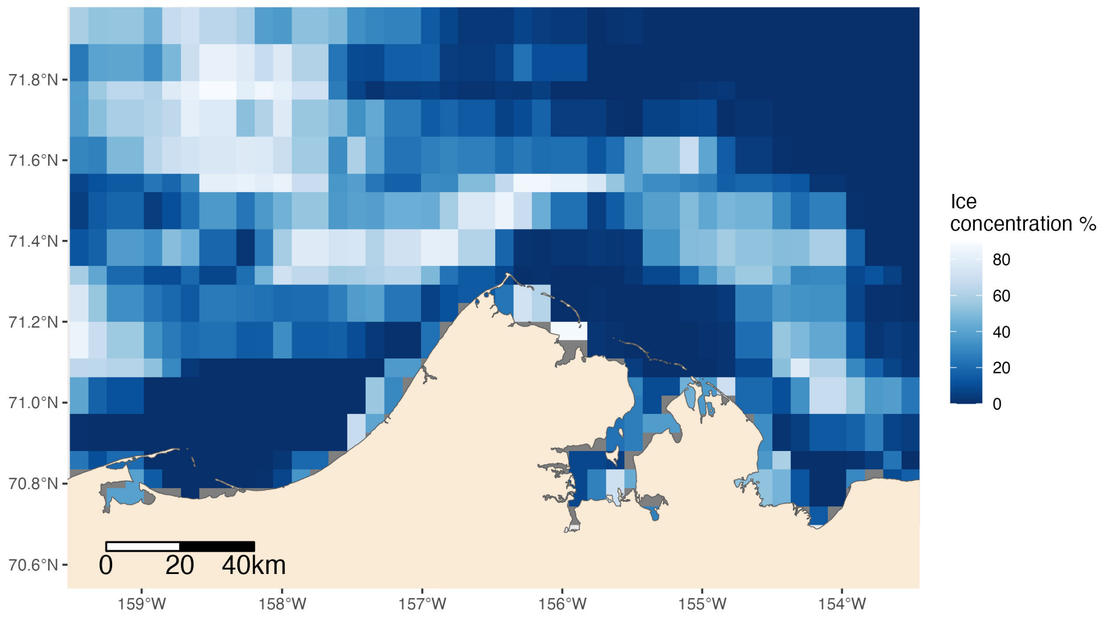


Figure S2: Ice concentration (%) July 20, 2016. Areas of dark blue indicate lower ice concentration when white indicates areas of high ice concentration.

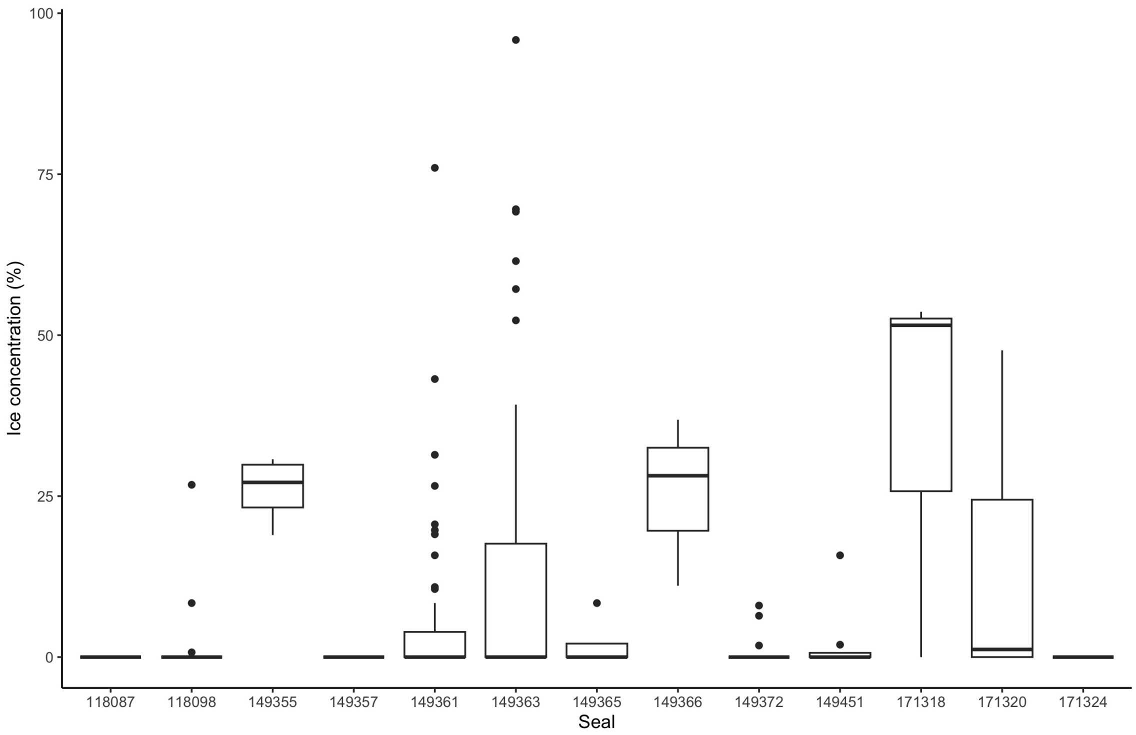


Figure S3: Ice concentration (%) at used locations for each ringed seal.
